# Protenix-v1: Toward High-Accuracy Open-Source Biomolecular Structure Prediction

**DOI:** 10.64898/2026.02.05.703733

**Authors:** Protenix Team, Yuxuan Zhang, Chengyue Gong, Hanyu Zhang, Wenzhi Ma, Zhenyu Liu, Xinshi Chen, Jiaqi Guan, Lan Wang, Yanping Yang, Yu Xia, Wenzhi Xiao

## Abstract

We introduce Protenix-v1 (PX-v1), the first fully open-source structure prediction model to attain superior performance to AlphaFold3 while strictly adhering to the same training data cutoff, model size, and inference budget. Beyond standard evaluations, we highlight the effectiveness of inference-time scaling behavior of Protenix-v1, demonstrating that increasing the sampling budget yields consistent improvements in prediction quality—a behavior previously observed in AlphaFold3 and largely absent from prior open-source models. In addition to improved accuracy, Protenix-v1 incorporates key capabilities including protein template integration and RNA MSA support. Furthermore, to better support real-world applications such as drug discovery, we additionally release Protenix-v1-20250630, a variant trained on a larger dataset (cutoff: June 30, 2025), delivering further improved prediction accuracy. Finally, we identify limitations in existing benchmarking practices and provide updated evaluation tools and year-stratified benchmarks to support more reliable and transparent assessment. Collectively, these contributions provide a robust foundation for the Protenix series and the broader field.

## 1 Introduction

Structure prediction models have become indispensable tools in both fundamental biological research [e.g., 10, 12, 25] and drug discovery [e.g., 6, 20, 24]. Although the open-source ecosystem has expanded rapidly [e.g. 5, 7, 18, 19, 26, 29], a performance gap persists between current open-source implementations and AlphaFold3 [23, 30]. Moreover, closed-source models, due to their non-transparent nature, impede systematic comparisons and thus create substantial barriers to a comprehensive and impartial assessment.

In this work, we introduce Protenix-v1, the first fully open-source biomolecular structure prediction model to reach or exceed AlphaFold3-level performance while maintaining the same training data cutoff, model scale, and inference budget. Under these controlled conditions, Protenix-v1 demonstrates robust performance across diverse benchmark sets, establishing that the long-standing performance gap between fully open-source models and AlphaFold3 is not fundamental.

A key observation of Protenix-v1 is its inference-time scaling behavior. For challenging targets, such as antibody–antigen complexes, increasing the sampling budget from a baseline level to hundreds of candidates yields consistent, approximately log-linear improvements in prediction accuracy. This property, previously observed in AlphaFold3 [1], has been largely absent from prior open-source models and suggests that Protenix-v1 operates within a similar performance regime. Importantly, this behavior provides users with a practical control knob, enabling explicit trade-offs between computational cost and prediction accuracy.

Beyond inference behavior, Protenix-v1 adopts enhanced data processing pipelines and incorporates additional input features such as RNA MSA support and protein template integration. These are accompanied by a more complete alignment with the training data components described in AlphaFold3, including expanded disorder-focused distillation and large-scale monomer distillation based on MGnify.

To better support practical applications, we additionally release Protenix-v1-20250630, a variant trained on a more recent and less restrictive dataset. While Protenix-v1 is designed for controlled comparison under strict data cutoffs, the expanded variant leverages additional structural data to improve performance on newly released targets commonly encountered in applied settings such as drug discovery. This dual-release strategy distinguishes benchmark-aligned evaluation from real-world deployment, allowing users to select the model variant most appropriate for their use case.

For model evaluation, we construct what is, to our knowledge, the largest available test set, which mitigates dataset bias by expanding the diversity of interface clusters and increasing the number of intra-cluster examples. We leverage bootstrapping to alleviate randomness-induced variance in data-scarce domains, such as antibody-antigen complexes. Using this evaluation framework, we systematically characterize the model’s performance across multiple tasks. Figure 1 and Figure 2 summarize these results. Figure 1 compares Protenix-v1 against existing models on the FoldBench and demonstrates inference-time scaling power on both FoldBench and PXM-22to25-Antibody, while Figure 2 summarizes the key results for the other proposed benchmark suite. In the following sections, we compare Protenix-v1 against its immediate predecessor, Protenix-v0.5.0, and other representative models under the same evaluation framework.

**Figure 1.**
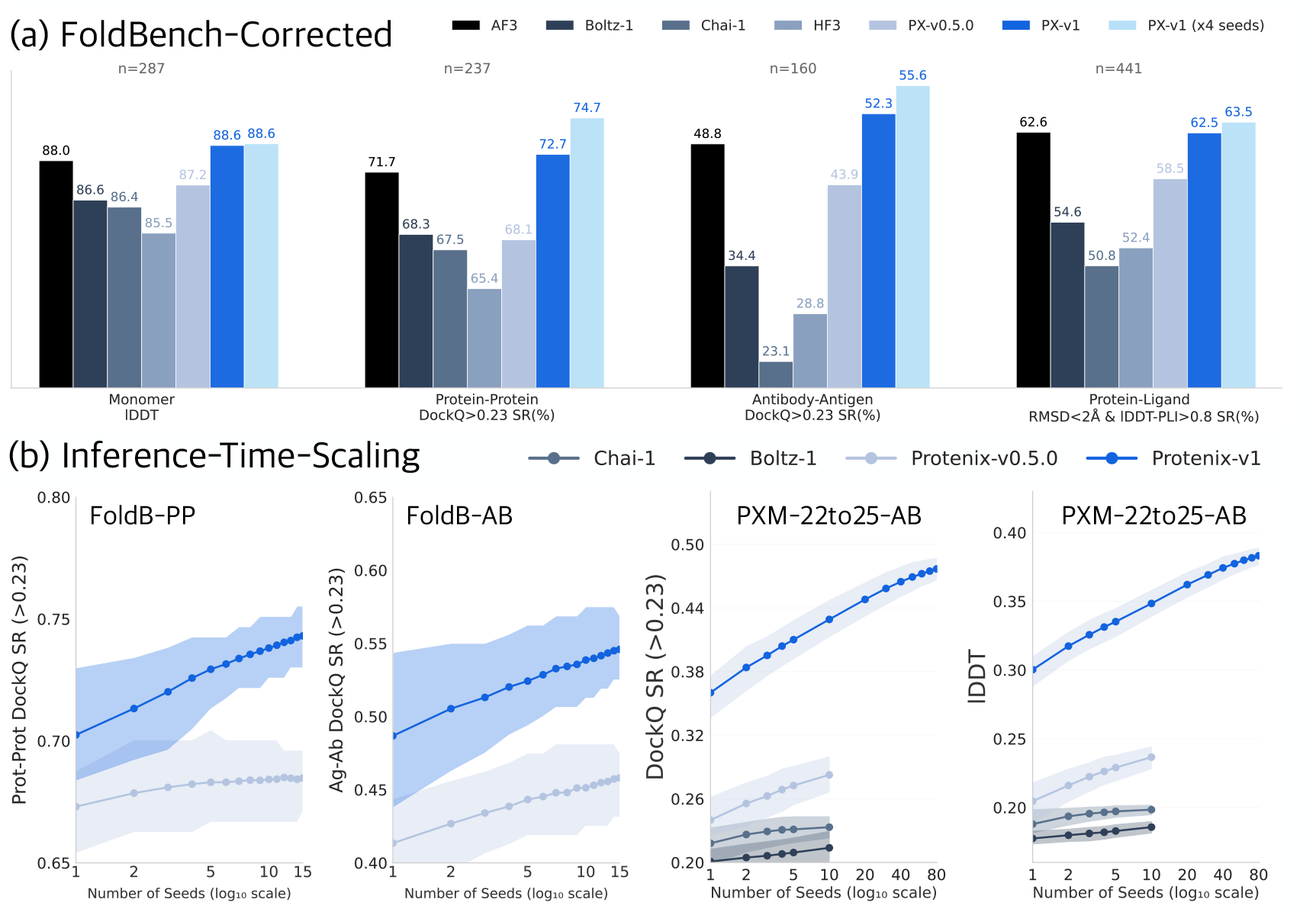
Evaluation Results. (a) We construct a comparable subset of FoldBench using official per-instance logs. Some FoldBench metrics reported by individual methods are computed on evaluation subsets that are not fully specified or lack variance reporting, making it difficult to reliably determine the underlying data coverage; these results are therefore omitted from the table. Protenix-v1 results are bootstrapped from 20 seeds, with aggregate results from all 20 seeds also reported. AlphaFold3 results are sourced from the official FoldBench report. ‘SR’ refers to success rate. (b) Protenix-v1 exhibits robust inference-time scaling on PXM-22to25-Antibody and FoldBench subsets. On PXM-22to25-Antibody, Protenix-v1 results are bootstrapped from 100 seeds.

**Figure 2.**
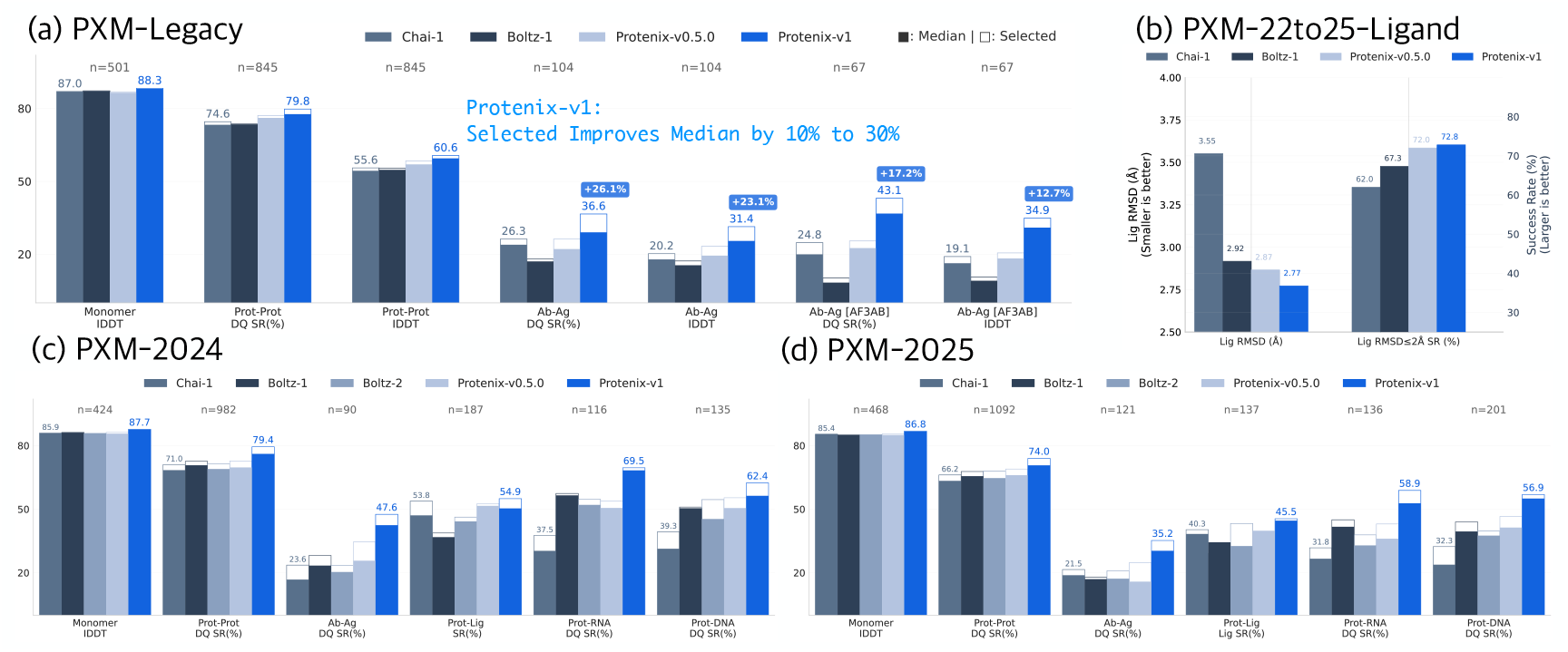
PXM Main Results. Performance across distinct PXM subsets are listed. ‘Median’ denotes the median performance across 5 × 5 sampled structures (solid filled bars), while ‘Selected’ denotes the performance of structures selected by confidence score (outlined bars with no fill).

## 2 Evaluation Setup

We evaluate Protenix-v1 using a comprehensive set of benchmarks, encompassing both established public datasets and our newly constructed, year-stratified evaluation suites. This section describes the benchmark composition, the protocols adopted to ensure fair and reproducible comparisons, and the inference configurations employed throughout our study.

### Evaluation Benchmarks

To ensure broad coverage across molecular modalities and prediction tasks, we evaluate Protenix-v1 on a diverse suite of benchmarks summarized in Table 1. These benchmarks fall into two categories. The first category comprises publicly available benchmarks, including Runs-N-Poses [23] for protein–ligand co-folding and FoldBench [30] for general biomolecular interaction prediction. For these two benchmarks, the evaluation results for all models other than Protenix-v0.5.0 and Protenix-v1 are collected from the publicly available repositories accompanying the original papers, without re-running those models. The second category consists of our newly curated, year-stratified PXM evaluation suites, where both dataset construction and metric computation follow PXMeter [13] and are implemented using the PXMeter v1.0.0 codebase^1^. Specifically, we introduce PXM-2024 and pxm-2025, which are curated from PDB entries released in 2024 and 2025, respectively. For data-scarce but practically important tasks such as protein–ligand and protein–antibody interface prediction, single-year datasets often lack sufficient statistical power. To address this, we additionally aggregate structures released between 2022 and 2025 to construct two task-specific benchmarks: PXM-22to25-Ligand and pxm-22to25-antibody. Apart from these newly curated benchmark sets, we also include the PXM-Legacy benchmark introduced in our prior work, which aligns with the evaluation window of AlphaFold3 (2022-05-01 to 2023-01-12). To specifically evaluate Protenix-v1-20250630 under a strict post-cutoff setting, we further curate pxm-2025h2, which comprises PDB entries released in the second half of 2025. Filtering criteria, redundancy control, and curation procedures for these datasets are provided in Appendix A.1.

**Table 1.** Benchmark Datasets. This table summarizes time cutoff window, total number of complexes, and targeted prediction domain for each dataset. The first group comprises existing open-access benchmarks, while the second presents distinct subsets of our evaluation benchmark, which features a substantially larger number of complexes.

| Test Set | Reference | Time Cutoff | #Complexes | Prediction Task |
| --- | --- | --- | --- | --- |
| RUNS-N-POSES | Škrinjar et al. [23] | 2021-09-30 to 2025-01-09 | 2600 | Ligands |
| FOLDBENCH | Xu et al. [30] | 2021-09-30 to 2024-11-01 | 1522 | All |
| PXM-LEGACY | Ma et al. [13] | 2022-05-01 to 2023-01-12 | 1567 | All |
| PXM-2024 | new | 2024-01-01 to 2024-12-31 | 2051 | All |
| PXM-2025 | new | 2025-01-01 to 2025-12-31 | 2359 | All |
| PXM-2025H2 | new | 2025-07-01 to 2025-12-31 | 1159 | All |
| PXM-22To25-LIGAND | new | 2022-01-01 to 2025-12-31 | 629 | Ligands |
| PXM-22To25-ANTIBODY | new | 2022-01-01 to 2025-12-31 | 528 | Protein-Antibody |

### Fair Evaluation Protocols

Unless otherwise specified, all evaluations use five random seeds, with each seed generating five diffusion samples. The number of recycles is fixed to 10 during inference. Final predictions are selected using the ranking protocol specified by each benchmark. For Runs-N-Poses, structures with higher <u>ranking score</u> are selected. For FoldBench, we follow the original benchmark ranking definition. For all other datasets, we use the <u>chain-pair ipTM score</u> for interface selection and the <u>chain pTM</u> for chain selection, consistent with the protocol of AlphaFold3. We mark the selected results using confidence scores as <u>‘Selected’</u>. Performance metrics are reported exclusively on the **common intersection** of successfully evaluated samples across compared models, avoiding discrepancies caused by model-specific inference or evaluation failures.

### Revisiting FoldBench: Data Coverage and Variance Considerations

Fold-Bench remains a valuable resource for evaluating general biomolecular structure prediction, particularly as one of the few benchmarks enabling direct comparison with AlphaFold3. However, careful examination reveals important limitations in data coverage and statistical variance that affect cross-model comparability. In FoldBench, model-specific inference or evaluation failures lead to inconsistent data coverage across methods, such that each model is effectively evaluated on a different subset of the targets. This behavior, also noted in the original FoldBench report, is particu-larly pronounced for large complexes, where less memory-efficient models often encounter Out-of-Memory (OOM) failures. Although the aggregated metrics are reported in the FoldBench Supplementary Table 3 and on the official leaderboard, and subsequently cited by other studies, the underlying evaluation procedure does not enforce a common intersection of samples when computing these metrics. As illustrated in Figure 3, aggregated metrics computed without enforcing a common intersection of successfully evaluated samples inherently reflect differences in evaluation subsets rather than model performance alone, producing misleading cross-model rankings. In addition, several FoldBench subsets contain a limited number of samples, making the reported metrics highly sensitive to stochastic variation. As shown in Figure 1b, a 20-random-seed bootstrapping analysis shows that the 95% confidence interval for Protenix-v1’s DockQ SR (based on a single run of “5 seeds × 5 samples” protocol) spans from 49.3% to 56.3%. This substantial fluctuation underscores two methodological requirements for reliable evaluation: sufficient test set scale to ensure statistical power, and explicit variance-aware evaluation protocols. Nevertheless, as FoldBench remains one of the few benchmarks facilitating a direct comparison with AlphaFold3, we utilize it in our evaluation, applying strict corrections to mitigate these biases.

**Figure 3.**
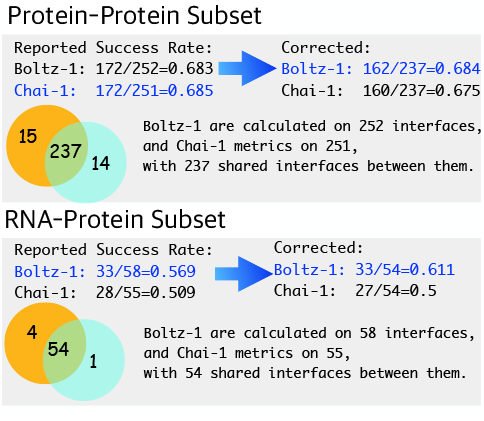
FoldBench Examples.

**Figure 4.**
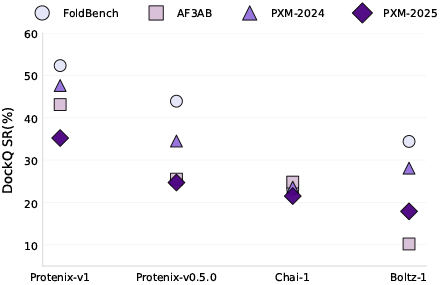
Antibody Test Sets. Existing antibody-antigen tests yield inconsistent conclusions.

### Correction of FoldBench for Fair Comparison

Motivated by the considerations above, we construct a corrected FoldBench subset by identifying the common intersection of successfully evaluated samples across all compared models and reporting performance on this shared set. In addition, Protenix-v1 is evaluated using a 20-seed bootstrapping protocol to improve the stability and reliability of the reported metrics.

### Inference Configuration of Protenix-1

For Protenix-v1, we adopt a diffusion-based inference configuration consistent with the hyperparameter settings of AlphaFold3. Variability across inference runs is introduced through the following randomness: the MSA input is subsampled to a maximum of *k* sequences, where *k* is sampled uniformly from the range [1, 16384]; inference-time dropout is applied to the pair embeddings; the stochastic sampling trajectory of the diffusion process also introduces randomness. Details of the MSA and Template search procedure are provided in Appendix A. For the evaluation of Protenix-v1, we utilize protein MSAs, RNA MSAs, and protein templates as input features. For the sake of fair comparison and ablation analysis, we additionally report the performance of Protenix-v1 using ablated configurations where RNA MSAs are excluded. The results under these settings are denoted as <u>Protenix-v1-wo-RNA-MSA</u>.

## 3 Protenix-v1 Achieves SOTA Across Modalities

To facilitate a comprehensive comparison, we evaluate Protenix-v1 using a diverse set of metrics, including DockQ [2], lDDT [14], and ligand-associated metrics [13]. For protein–ligand tasks, the PB-Valid success rate is defined as the percentage of predictions satisfying all 18 validity criteria in PoseBusters [3] used by AlphaFold3.

### FoldBench

Table 2 summarizes the performance on the FoldBench benchmark under multiple evaluation settings. In the first panel, which reports results on the common intersection of successfully evaluated samples across all models, both AlphaFold3 and Protenix-v1 outperform existing open-source baselines across all evaluated domains. The second panel presents a direct comparison between Protenix-v1 and the official AlphaFold3 FoldBench results,restricted to the intersection of their respective inference outputs. While AlphaFold3 retains an advantage on protein–ligand and protein–DNA docking tasks, Protenix-v1 achieves higher accuracy on protein–protein and antibody–antigen (Ab–Ag) interface prediction. For completeness, the third panel lists the original FoldBench metrics as reported in the official benchmark. We emphasize that these values are not directly comparable across models due to inconsistent evaluation subsets.

**Table 2.** FoldBench Results. The first panel presents the common intersection of all models available in FoldBench, with subsets determined using the official files provided by the benchmark authors. The second panel displays the intersection between Protenix-v1 inference outputs and the official AlphaFold3 results provided by FoldBench. The third panel lists the original metrics reported by FoldBench for reference, where values in parentheses indicate the size of the data subsets used for calculation. The ^old^ superscript refers to the performance of Protenix-v0.2.0 as reported in FoldBench. Protenix-v1 results are generated via bootstrapping 5 seeds from 20 random seeds (seeds 101–120), while Protenix-v1 (×4 seeds) apply all 20 seeds. For each panel, the first row (“# chains /interfaces”) indicates the number of samples remaining after intersection. Some FoldBench metrics reported by individual methods are computed on evaluation subsets that are not fully specified or lack variance reporting, making it difficult to reliably determine the underlying data coverage; these results are therefore omitted from the table.

| Domain | Monomer | Prot-Prot | Ab-Ag | Prot-Lig | Prot-RNA | Prot-DNA | DNA | RNA |
| --- | --- | --- | --- | --- | --- | --- | --- | --- |
| Metric | IDDT | DockQ SR(%) | DockQ SR(%) | SR(%) | DockQ SR(%) | DockQ SR(%) | IDDT | IDDT |
| # chains / interfaces | 287 | 237 | 160 | 441 | 46 | 274 | 14 | 15 |
| AlphaFold3 | 0.8803 | 71.73 | 48.75 | 62.59 | 65.22 | **75.91** | 0.5278 | 0.6140 |
| Boltz-1 | 0.8664 | 68.35 | 34.38 | 54.65 | 65.22 | 67.15 | 0.3388 | 0.4416 |
| Chai-1 | 0.8639 | 67.51 | 23.12 | 50.79 | 50.00 | 66.06 | 0.4574 | 0.4852 |
| HF3 | 0.8547 | 65.40 | 28.75 | 52.38 | 52.17 | 49.27 | 0.2909 | 0.5487 |
| Protenix-v0.2.0<sup>old</sup> | 0.8622 | 66.24 | 35.62 | 51.02 | 56.52 | 64.60 | 0.4384 | 0.5903 |
| Protenix-v0.5.0 | 0.8717±0.0011 | 68.06±0.93 | 43.94±1.43 | 58.49±0.56 | 59.37±1.12 | 68.02±0.73 | 0.4960±0.0201 | 0.6046±0.0126 |
| Protenix-v1 | 0.8857±0.0007 | 72.70±0.76 | 52.31±1.43 | 62.54±0.73 | 68.46±1.43 | 69.13±0.76 | **0.5693**±0.0109 | **0.6547**±0.0088 |
| Protenix-v1 (×4 seeds) | **0.8863** | **74.68** | **55.63** | **63.49** | **69.57** | **69.71** | **0.5682** | **0.6523** |
| # chains / interfaces | 334 | 266 | 167 | 547 | 69 | 317 | 14 | 15 |
| AlphaFold3 | 0.8843 | 72.93 | 47.90 | **64.90** | 62.32 | **79.18** | 0.5278 | 0.6140 |
| Protenix-v0.5.0 | 0.8756±0.0008 | 69.55±0.99 | 42.23±1.11 | 59.45±0.60 | 51.49±1.84 | 72.34±0.65 | 0.4986±0.0187 | 0.6053±0.0137 |
| Protenix-v1 | 0.8874±0.0006 | 74.00±0.72 | 50.12±1.36 | 62.79±0.69 | 66.67±1.29 | 73.29±0.61 | **0.5693**±0.0109 | **0.6547**±0.0088 |
| Protenix-v1 (×4 seeds) | **0.8881** | **75.94** | **53.29** | **63.99** | **68.12** | **73.82** | **0.5682** | **0.6523** |
| # chains / interfaces | **Incomparable:** FOLDBENCH official metrics and their computed data sizes are included for reference. |  |  |  |  |  |  |  |
| AlphaFold3 | 0.8843(334) | 72.93(266) | 47.90(167) | 64.90(547) | 62.32(69) | 79.18(317) | 0.5278(14) | 0.6140(15) |
| Chai-1 | 0.8664(322) | 68.53(251) | 23.64(165) | 51.23(476) | 50.91(55) | 69.97(313) | 0.4574(14) | 0.4852(15) |
| Boltz-1 | 0.8694(330) | 68.25(252) | 33.54(164) | 55.04(476) | 56.90(58) | 70.97(310) | 0.3388(14) | 0.4416(15) |

### PXM-2024 and PXM-2025

Results on the year-stratified PXM-2024 and PXM-2025 benchmarks are reported in Table 3. Across all six evaluated domains and metrics, Protenix-v1 consistently outperforms representative open-source baselines. Notably, Protenix-v1 demonstrates strong gains on protein–protein interface prediction tasks, achieves an approximately 10% relative improvement over current leading open-source baselines on PXM-2024. Additionally, we find that the confidence head of Protenix-v1 effectively facilitates the selection of higher-quality structures. As illustrated in Figure 2, confidence-based selection consistently elevates prediction quality relative to the median across all domains, with particularly large margins observed for antibody–antigen and protein–DNA interfaces.

**Table 3.** PXM-2024 and PXM-2025 Results. In the first and second panel, we list the PXM-2024 and PXM-2025 results, respectively. ‘DQ’ denotes DockQ, SR denotes success rate, ‘RMSD SR’ refers to the rate of ligand RMSD<2Å, and ‘Lig SR’ refers to both RMSD *<* 2Å and PB-Valid. More detailed ligand metrics are reported on PXM-22to25-ligands.

| Domain | Monomer | Prot-Prot(non-ab) |  | Ab-Ag |  | Prot-Lig |  | Prot-RNA |  | Prot-DNA |  |
| --- | --- | --- | --- | --- | --- | --- | --- | --- | --- | --- | --- |
| # Complex / # Cluster | 424/280 | 982/745 |  | 90/78 |  | 187/74 |  | 116/84 |  | 135/176 |  |
| Metric | IDDT | DQ SR(%) | IDDT | DQ SR(%) | IDDT | RMSD SR(%) | Lig SR(%) | DQ SR(%) | IDDT | DQ SR(%) | IDDT |
| Chai-1 | 0.8592 | 71.03 | 0.5262 | 23.59 | 0.2330 | 65.59 | 53.79 | 37.54 | 0.2380 | 39.29 | 0.2765 |
| Boltz-1 | 0.8622 | 72.58 | 0.5358 | 28.13 | 0.2323 | 73.08 | 38.75 | 57.40 | 0.3302 | 50.99 | 0.3649 |
| Boltz-2 | 0.8568 | 71.29 | 0.5281 | 23.44 | 0.1929 | 68.60 | 46.25 | 54.63 | 0.3592 | 54.51 | 0.3487 |
| Protenix-v0.5.0 | 0.8622 | 72.75 | 0.5505 | 34.53 | 0.2788 | 71.31 | 52.55 | 53.85 | 0.3259 | 55.39 | 0.3878 |
| Protenix-v1 | **0.8770** | **79.42** | **0.6003** | **47.56** | **0.3988** | **77.13** | **54.95** | **69.53** | **0.4170** | **62.38** | **0.4214** |
| # Complex / # Cluster | 468/240 | 1092/821 |  | 121/87 |  | 137/94 |  | 136/94 |  | 201/267 |  |
| Metric | IDDT | DQ SR(%) | IDDT | DQ SR(%) | IDDT | RMSD SR(%) | Lig SR(%) | DQ SR(%) | IDDT | DQ SR(%) | IDDT |
| Chai-1 | 0.8542 | 66.15 | 0.4922 | 21.51 | 0.1761 | 54.48 | 40.32 | 31.76 | 0.2422 | 32.33 | 0.2441 |
| Boltz-1 | 0.8505 | 67.77 | 0.5090 | 17.93 | 0.1832 | 61.78 | 29.00 | 44.88 | 0.2879 | 43.99 | 0.3024 |
| Boltz-2 | 0.8508 | 67.79 | 0.5093 | 20.94 | 0.1724 | 57.58 | 43.21 | 37.87 | 0.2487 | 39.72 | 0.2830 |
| Protenix-v0.5.0 | 0.8558 | 68.94 | 0.5300 | 24.74 | 0.2123 | **70.56** | 42.67 | 43.01 | 0.2922 | 46.57 | 0.3248 |
| Protenix-v1 | **0.8682** | **73.96** | **0.5617** | **35.22** | **0.3074** | 70.20 | **45.49** | **58.85** | **0.3717** | **56.88** | **0.3945** |

### PXM-Legacy

We report the performance on the 2022–2023 PXM-Legacy benchmark in Table 4. Protenix-v1 achieves higher median performance than all existing open-source baselines. Moreover, as shown in Figure 2a, confidence-based selection substantially improves upon the median prediction quality. Especially, for antibody-antigen complexes, selected predictions of Protenix-v1 yield gains of 10% to 30% in both DockQ SR and lDDT, demonstrating the effectiveness of the confidence head in identifying high-quality structures from the sampled ensemble.

**Table 4.** PXM-Legacy Results. Protenix-v1 improves Protenix-v0.5.0 by a large margin. For the AF3AB benchmark, we adopt the interface clustering results provided in the AlphaFold3 metadata to ensure alignment between the clustering of evaluation sets. We report the metrics calculated on the intersection of all models’ outputs, as Boltz-1 and Chai-1 failed for certain cases. We extract the AlphaFold3 results from the original report.

| Domain | Monomer | Prot-Prot(non-ab) |  | Ab-Ag |  | Ab-Ag [AF3AB] |  |
| --- | --- | --- | --- | --- | --- | --- | --- |
| # Complex / # Cluster | 501/309 | 845/695 |  | 104/58 |  | 67/61 |  |
| Metric | IDDT | DQ SR(%) | IDDT | DQ SR(%) | IDDT | DQ SR(%) | IDDT |
| AlphaFold3 | - | - | - | - | - | **44** | - |
| Boltz-1 | 0.8730 | 73.69 | 0.5545 | 18.02 | 0.1723 | 10.21 | 0.1056 |
| Chai-1 | 0.8701 | 74.57 | 0.5556 | 26.28 | 0.2023 | 24.77 | 0.1905 |
| Protenix-v0.5.0 | 0.8687 | 77.14 | 0.5847 | 26.25 | 0.2326 | 25.47 | 0.2055 |
| Protenix-v1 | **0.8832** | **79.78** | **0.6064** | **36.58** | **0.3136** | **43.07** | **0.3488** |

### PXM-22to25-Antibody

Given that conventional antibody datasets in FoldBench or AF3AB typically contain very limited cluster diversity, which can introduce evaluation bias or high variance, we construct a dedicated antibody benchmark featuring a significantly larger number of targets. Furthermore, to ensure statistically robust results and mitigate stochasticity, we conduct inference across 20 random seeds for this evaluation suite.

We then report the bootstrapped mean and 95% confidence intervals (CI) for the standard “5 seeds × 5 samples” protocol by resampling from this 20-seed (101–120) pool.

As shown in Table 5, Protenix-v1 achieves superior median, selected, and best performance across most metrics compared to existing baselines, with the exception of Boltz-2, whose results are partially confounded by training data overlap.

**Table 5.** PXM-22to25-Antibody and PXM-22to25-Ligand Results. ^*^ denotes that this evaluation dataset contains part of Boltz-2 training data. antibody subset results are averaged across 498 complexes with 359 clusters, while ligand subset results are averaged across 625 complexes with 252 clusters, after intersection with Boltz-1 and Chai-1.

|  | DockQ SR(%) |  |  | IDDT |  |  | Lig RMSD (Å) | Lig RMSD<2Å SR(%) |
| --- | --- | --- | --- | --- | --- | --- | --- | --- |
|  | best | selected | median | best | selected | median | selected | selected |
| Boltz-2* | 50.35±0.67 | 40.10±0.31 | 38.59±0.48 | 0.4167±0.0037 | 0.3474±0.0028 | 0.3379±0.0014 | 2.939 | 72.95 |
| Chai-1 | 31.54±1.08 | 22.12±0.63 | 17.77±0.33 | 0.2535±0.0053 | 0.1911±0.0022 | 0.1630±0.0024 | 3.551 | 61.96 |
| Boltz-1 | 29.52±0.85 | 20.59±0.71 | 17.27±0.48 | 0.2326±0.0036 | 0.1840±0.0024 | 0.1877±0.0043 | 2.917 | 67.31 |
| Protenix-v0.5.0 | 44.03±1.86 | 27.50±0.96 | 21.21±0.70 | 0.3345±0.0093 | 0.2313±0.0048 | 0.1930±0.0045 | 2.866 | 72.01 |
| Protenix-v1 | **54.22±1.50** | **40.26±0.78** | **33.14±0.81** | **0.4130±0.0081** | **0.3310±0.0049** | **0.2842±0.0038** | **2.772** | **72.79** |

### PXM-22to25-Ligand

As demonstrated in Table 5, among the evaluated models, Protenix-v1 attains superior performance, achieving the lowest ligand RMSD (**2.772** Å) and the highest RMSD<2Å success rate (**72.79%**).

### Runs-N-Poses

To assess protein–ligand co-folding performance, we evaluate Protenix-v1 on the Runs-N-Poses test set. Similar to FoldBench, the publicly released results exhibit data incompleteness and subset mismatch across models. We therefore remove entries with fewer than 25 predictions (5 seeds × 5 diffusion samples) and restrict evaluation to the subset consistently reported by all compared methods. Under this controlled setting, Protenix-v1 improves upon Protenix-v0.5.0, as shown in Figure 5.

**Figure 5.**
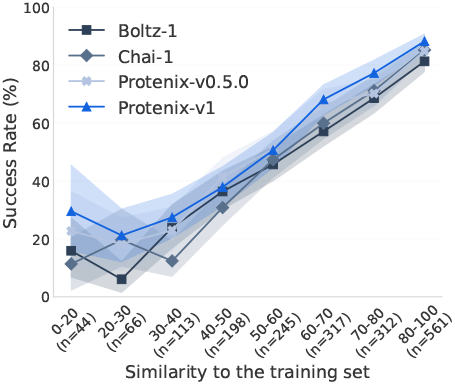
Runs-N-Poses.

### Inference Time Scaling Unlocks Performance via Computational Budget

On the PXM-22to25-Antibody benchmark, we demonstrate that Protenix-v1 exhibits enhanced performance as computational resources increase (e.g., utilizing up to 100 random seeds per complex). As illustrated in Figure 1b, Protenix-v1 achieves higher lDDT scores and DockQ Success Rates (SR) through additional inference-time sampling. Specifically, by bootstrapping from a pool of 100 seeds, we observe significant gains: the DockQ SR improves from 36.01% with a single seed to 42.92% with 10 seeds, reaching 47.68% with 80 seeds. These findings suggest a clear pathway for further optimization; we identify the distillation of these high-quality sampling signals into the base model as a primary direction for future work.

### A More Effective Design Filter

Confidence scores—such as ipTM, pTM, and pLDDT—are standard rankers for filtering candidates in protein binder and antibody design [e.g., 15, 20, 24]. Utilizing the empirical binder data released by Cao et al. [4], we validate that Protenix-v1 serves as a superior filter for identifying successful designs. We frame this evaluation as a binary classification task: distinguishing binding from non-binding designs for each target. For our test set, we include all provided positive examples, while negative examples are subsampled to a maximum of 20,000 per target to manage computational overhead.

As summarized in Table 6, Protenix-v1 achieves the highest Area Under the Curve (AUC) scores across nearly all targets. Baseline models, specifically Chai-1 and Boltz-1, notably outperform Protenix-v1 on only a single target: SC2RBD. Furthermore, in terms of Average Precision (AP), Protenix-v1 consistently delivers the leading performance for the majority of evaluated cases. While these results demonstrate the robustness of current confidence heads, exploring multi-score combinations and developing specialized filtering architectures are reserved for future investigation.

**Table 6.** Filter Results. We report the filter ability on open-source mini-binder design dataset with different models. Protenix-v1 gets the highest AUC scores and average precisions on most of the target.

| Metric |  | AUC |  |  |  |  |  |  |  | Average Precision |  |  |  |  |  |  |  |
| --- | --- | --- | --- | --- | --- | --- | --- | --- | --- | --- | --- | --- | --- | --- | --- | --- | --- |
| Target |  | FGFR2 | IL-7RA | InsulinR | PDGFR | SC2RBD | TGFb | TrkA | VirB8 | FGFR2 | IL-7RA | InsulinR | PDGFR | SC2RBD | TGFb | TrkA | VirB8 |
| Best {pTM, iPTM, pLDDT} | AF2-IG | 0.640 | 0.785 | 0.889 | **0.699** | 0.585 | 0.640 | 0.850 | 0.588 | 0.069 | 0.018 | **0.266** | **0.049** | **0.054** | 0.013 | 0.011 | 0.006 |
|  | Chai-1 | 0.706 | 0.854 | 0.877 | 0.691 | **0.748** | 0.662 | 0.774 | 0.581 | 0.098 | **0.088** | 0.189 | 0.034 | 0.031 | 0.030 | 0.008 | 0.008 |
|  | Boltz-1 | 0.722 | **0.901** | 0.902 | 0.670 | 0.698 | 0.688 | 0.820 | 0.605 | 0.102 | 0.033 | 0.143 | 0.035 | 0.008 | 0.033 | 0.069 | 0.009 |
| pTM | Protenix-v0.5.0 | 0.707 | 0.854 | 0.890 | 0.661 | 0.633 | 0.745 | 0.872 | 0.525 | 0.098 | 0.033 | 0.136 | 0.032 | 0.015 | 0.029 | 0.115 | 0.009 |
|  | Protenix-Mini | 0.713 | 0.698 | 0.863 | 0.668 | 0.646 | 0.624 | 0.892 | 0.563 | **0.134** | 0.005 | 0.133 | 0.042 | 0.011 | 0.026 | **0.063** | 0.007 |
|  | Protenix-v1 | 0.725 | 0.878 | 0.926 | 0.655 | 0.704 | 0.777 | **0.903** | 0.580 | 0.121 | 0.022 | 0.134 | 0.029 | 0.029 | 0.050 | **0.151** | 0.010 |
| ipTM | Protenix-v0.5.0 | 0.703 | 0.835 | 0.891 | 0.677 | 0.622 | 0.749 | 0.847 | 0.552 | 0.100 | 0.043 | 0.135 | 0.048 | 0.012 | 0.031 | 0.109 | **0.013** |
|  | Protenix-Mini | 0.712 | 0.770 | 0.860 | 0.681 | 0.472 | 0.636 | 0.864 | 0.575 | **0.134** | 0.012 | 0.139 | 0.046 | 0.006 | 0.026 | 0.104 | 0.007 |
|  | Protenix-v1 | 0.724 | 0.872 | **0.929** | 0.696 | 0.695 | 0.782 | 0.887 | 0.603 | 0.129 | 0.025 | 0.141 | **0.049** | 0.023 | **0.053** | **0.149** | **0.012** |
| pLDDT | Protenix-v0.5.0 | 0.729 | 0.871 | 0.894 | 0.687 | 0.645 | 0.778 | 0.857 | 0.548 | 0.100 | **0.073** | 0.136 | 0.032 | 0.004 | 0.032 | 0.049 | 0.010 |
|  | Protenix-Mini | 0.729 | 0.690 | 0.860 | 0.684 | 0.700 | 0.656 | 0.873 | 0.599 | 0.131 | 0.005 | 0.121 | 0.038 | 0.007 | 0.020 | 0.023 | 0.008 |
|  | Protenix-v1 | **0.738** | **0.886** | 0.925 | **0.697** | **0.706** | **0.791** | 0.897 | **0.633** | 0.122 | 0.033 | 0.127 | 0.032 | 0.039 | 0.050 | 0.012 | **0.012** |

**Table 7.** PXM-2024 Ablation Results. Protenix-v1-wo-RNA-MSA omits RNA MSA inputs compared to Protenix-v1.

| Domain | Monomer | Prot-Prot(non-ab) |  | Ab-Ag |  | Prot-Lig |  | Prot-RNA |  | Prot-DNA |  |
| --- | --- | --- | --- | --- | --- | --- | --- | --- | --- | --- | --- |
| Metric | IDDT | DQ SR(%) | IDDT | DQ SR(%) | IDDT | RMSD SR(%) | Lig SR(%) | DQ SR(%) | IDDT | DQ SR(%) | IDDT |
| # Complex / # Cluster | 424/280 | 982/745 |  | 90/78 |  | 186/73 |  | 116/84 |  | 135/176 |  |
| Protenix-v1 | 0.8770 | 79.42 | 0.6003 | 47.56 | 0.3988 | 77.13 | 55.70 | 69.53 | 0.4170 | 62.38 | 0.4214 |
| Protenix-v1-wo-RNA-MSA | 0.8772 | 78.73 | 0.5981 | 51.41 | 0.3835 | 77.13 | 55.70 | 64.27 | 0.3874 | 59.90 | 0.4189 |

**Table 8.** FoldBench Ablation Results. Protenix-v1-wo-RNA-MSA omits RNA MSA inputs and is bootstrapped with 20 random seeds (101–120).

| Domain | Monomer | Prot-Prot | Ab-Ag | Prot-Lig | Prot-RNA | Prot-DNA | DNA | RNA |
| --- | --- | --- | --- | --- | --- | --- | --- | --- |
| Metric | IDDT | DockQ SR(%) | DockQ SR(%) | SR(%) | DockQ SR(%) | DockQ SR(%) | IDDT | IDDT |
| # chains / interfaces | 334 | 266 | 167 | 547 | 69 | 317 | 14 | 15 |
| AlphaFold3 | 0.8843 | 72.93 | 47.90 | 64.90 | 62.32 | 79.18 | 0.5278 | 0.6140 |
| Protenix-v1 | 0.8874±0.0006 | 74.00±0.72 | 50.12±1.36 | 62.79±0.69 | 66.67±1.29 | 73.29±0.61 | 0.5693±0.0109 | 0.6547±0.0088 |
| Protenix-v1-wo-RNA-MSA | 0.8875±0.0007 | 74.12±0.71 | 49.59±1.37 | 62.65±0.68 | 65.57±1.47 | 72.69±0.58 | 0.5771±0.0065 | 0.6107±0.0074 |

## 4 Ablation Studies of More Different Protenix Versions

### Protenix-v1-wo-RNA-MSA Results

We extend the comparison by evaluating Protenix-v1 against its variants that exclude RNA MSA inputs. As displayed in the below tables, without RNA MSA, protein-RNA interface and RNA monomer results drop, while the other domain gets quite similar results.

### Protenix-v1-20250630 Results

While Protenix-v1 keeps the same training data time-cutoff as AlphaFold3, we additionally release Protenix-v1-20250630, a variant trained on an expanded dataset for practical real-world applications. Protenix-v1-20250630 is trained on a dataset with time cutoff June 30th, 2025. The test set pxm-2025h2 constitutes a low-homology subset relative to the model’s training data. We report the results in Table 9. First, Protenix-v1-20250630 achieves superior performance on PXM-2024, a dataset included in the model’s training corpus. While this does not reflect generalization ability, it offers significant value for practical applications like drug discovery, where targets frequently share homology with recently released PDB structures. Second, on the pxm-2025h2 dataset, it yields comparable results on most domains, while significantly outperforming Protenix-v1 on Antibody-Antigen task, underscoring the value of data scaling in data-sparse regimes.

**Table 9.**
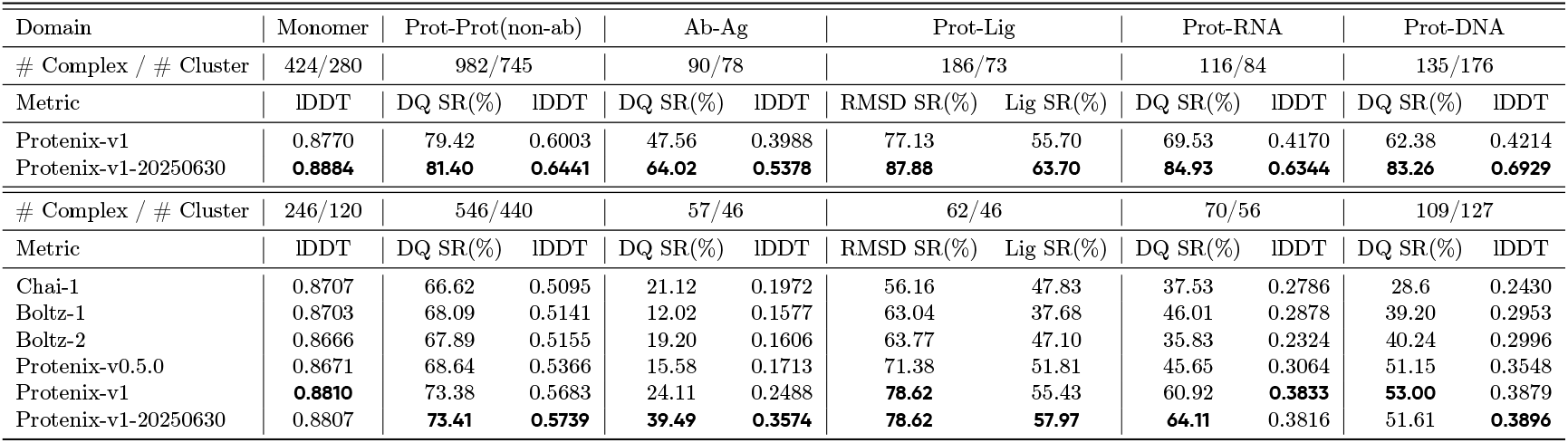
Protenix-v1-20250630 Ablation. We demonstrate the results of Protenix-v1-20250630 on PXM-2024 (part of full model’s training data) and PXM-2025H2 (Protenix-v1-20250630 test set).

| Domain | Monomer | Prot-Prot(non-ab) |  | Ab-Ag |  | Prot-Lig |  | Prot-RNA |  | Prot-DNA |  |
| --- | --- | --- | --- | --- | --- | --- | --- | --- | --- | --- | --- |
| # Complex / # Cluster | 424/280 | 982/745 |  | 90/78 |  | 186/73 |  | 116/84 |  | 135/176 |  |
| Metric | IDDT | DQ SR(%) | IDDT | DQ SR(%) | IDDT | RMSD SR(%) | Lig SR(%) | DQ SR(%) | IDDT | DQ SR(%) | IDDT |
| Protenix-v1 | 0.8770 | 79.42 | 0.6003 | 47.56 | 0.3988 | 77.13 | 55.70 | 69.53 | 0.4170 | 62.38 | 0.4214 |
| Protenix-v1-20250630 | **0.8884** | **81.40** | **0.6441** | **64.02** | **0.5378** | **87.88** | **63.70** | **84.93** | **0.6344** | **83.26** | **0.6929** |
| # Complex / # Cluster | 246/120 | 546/440 |  | 57/46 |  | 62/46 |  | 70/56 |  | 109/127 |  |
| Metric | IDDT | DQ SR(%) | IDDT | DQ SR(%) | IDDT | RMSD SR(%) | Lig SR(%) | DQ SR(%) | IDDT | DQ SR(%) | IDDT |
| Chai-1 | 0.8707 | 66.62 | 0.5095 | 21.12 | 0.1972 | 56.16 | 47.83 | 37.53 | 0.2786 | 28.6 | 0.2430 |
| Boltz-1 | 0.8703 | 68.09 | 0.5141 | 12.02 | 0.1577 | 63.04 | 37.68 | 46.01 | 0.2878 | 39.20 | 0.2953 |
| Boltz-2 | 0.8666 | 67.89 | 0.5155 | 19.20 | 0.1606 | 63.77 | 47.10 | 35.83 | 0.2324 | 40.24 | 0.2996 |
| Protenix-v0.5.0 | 0.8671 | 68.64 | 0.5366 | 15.58 | 0.1713 | 71.38 | 51.81 | 45.65 | 0.3064 | 51.15 | 0.3548 |
| Protenix-v1 | **0.8810** | 73.38 | 0.5683 | 24.11 | 0.2488 | **78.62** | 55.43 | 60.92 | **0.3833** | **53.00** | 0.3879 |
| Protenix-v1-20250630 | 0.8807 | **73.41** | **0.5739** | **39.49** | **0.3574** | **78.62** | **57.97** | **64.11** | 0.3816 | 51.61 | **0.3896** |

## 5 Conclusion

In this work, we present Protenix-v1, an open-source biomolecular structure prediction model that operates at the performance level of AlphaFold3 under matched training data cutoffs, model scale, and inference budgets. Through comprehensive evaluation across multiple domains, we demonstrate that Protenix-v1 consistently improves upon prior open-source models and exhibits stable behavior across a diverse set of biomolecular tasks. Beyond aggregate accuracy, we systematically characterize the inference-time scaling behavior of Protenix-v1, showing that increasing the sampling budget leads to consistent improvements in prediction quality on challenging targets such as antibody–antigen complexes. We further highlight limitations in existing evaluation benchmarks and propose principled corrections and variance-aware protocols to enable fair and reliable cross-model comparison. Alongside the release of Protenix-v1, we provide updated evaluation tools and expanded benchmark suites to facilitate transparent and reproducible assessment within the community.

Together, these contributions establish Protenix-v1 as a strong and practical foundation for open-source biomolecular structure prediction and provide a reference framework for future work on scalable inference, evaluation methodology, and real-world deployment.

## Acknowledgement

We thank Milong Ren, Jinyuan Sun, Zhaolong Li, Qinru Bai, Cong Liu, Xingang Peng, Song Xue, Jintao Zhu, Jiahui Tong and Qixu Cai for their careful review and valuable comments on the manuscript and case studies.

## Contributions

### Core Contributors

Yuxuan Zhang, Chengyue Gong, Hanyu Zhang, Wenzhi Ma, Zhenyu Liu, Xinshi Chen, Jiaqi Guan

### Contributors (Engineering Support)

Lan Wang, Yanping Yang, Yu Xia

### Team Lead

Wenzhi Xiao

## A Methods

### Training Data

Protenix-v1 is trained on a diverse corpus consisting of the following:

- **PDB Structures:** For Protenix-v1 we utilize a training cutoff of 2021-09-30, resulting in approximately 150k curated structures.
- **Monomer Distillation:** We construct a monomer distillation dataset based on the clustered MGnify database [17] (version 2019-05). Clusters containing fewer than 10 sequences are discarded. For each remaining cluster representative, we generate predictions using AlphaFold2 with its five original parameter sets, retaining the structure with the highest pLDDT. This process yields approximately 13 million distilled monomer structures. Notably, no additional filters regarding sequence length or pLDDT scores are applied. This updated dataset resolves the issue of low-quality structures present in our previous distillation corpus.
- **PDB Disorder Distillation:** Following the procedure described in AlphaFold3, we generate roughly 14K structures specifically for disorder distillation to improve robustness in intrinsically disordered regions. We empirically observe that incorporating this dataset effectively mitigates structural hallucination in disordered regions.

### Data Pipeline

Consistent with Protenix Team et al. [19], protein Multiple Sequence Alignments (MSAs) are generated using the colabfold pipeline [16]. We employ the UniRef [27] database for MSA pairing, utilizing taxonomy IDs for species identification and MSA pairing. While AlphaFold3 derives paired MSAs from UniProt and aggregates unpaired MSAs from diverse sources including UniRef, BFD, and MGnify, our colabfold-based pipeline operates primarily on uniref30 and colabfold_envdb. To maximize information retention, we retain UniRef sequences that fail the pairing process and incorporate them into the unpaired MSA features.

A significant addition in Protenix-v1 is the integration of templates and RNA MSAs:

- **Template Processing:** We adopt the template utilization strategy from AlphaFold3, capping the model at four templates. During inference, we select the top four templates released prior to 2021-09-30, ranked by *e*-value. During training, we randomly sample up to four templates released at least 60 days before the target sample’s release date to prevent data leakage. In addition to improving accuracy, we observe that template features lead to more stable model activations during training.
- **RNA MSA:** RNA MSA searching follows the AlphaFold3 pipeline. We utilize mmseqs2 for clustering and `nhmmer` [28] for searching across Rfam [9], RNAcentral [21], and the Nucleotide collection [22].

### Training Strategy

The training procedure for Protenix-v1 largely follows our previous framework [19]. Protenix-v1 are trained using multiple stages with increasing crop sizes of 384, 640, and 768, respectively. All training stages are conducted on a cluster of 256 GPUs. We train the confidence head together with the structure predictor.

### A.1 Additional details on test set generation

The three general test sets, PXM-2024, pxm-2025 and pxm-2025h2 are constructed primarily in accordance with the protocol established in Ma et al. [13], albeit without additional structural refinement. Supplementary to these comprehensive benchmarks, we curate two task-specific datasets: PXM-22to25-Ligand for the protein-ligand co-folding task and pxm-22to25-antibody for the antibody-antigen interface prediction task. Our curation pipeline first leverages PXMeter to generate a foundational benchmark set with a release date window from 2022-01-01 to 2025-12-31. Subsequently, we apply task-specific filters to extract the protein-ligand and antibody-antigen subsets.

#### A.1.1 Curation of Protein-Ligand Evaluation Benchmarks

To ensure a high-fidelity evaluation of protein-ligand co-folding, we implement a hierarchical filtering pipeline for ligand chains (excluding glycans and ions). The curation process is divided into three stages: basic structural filtering, chemical redundancy removal, and advanced quality validation.

##### Basic Structural Criteria

We first select ligand chains based on the following rigorous criteria to ensure experimental reliability and simplicity:

- The PDB entry must be determined uniquely via “X-RAY DIFFRACTION”.
- The experimental resolution must be ≤ 2.0 Å.
- All atoms within the ligand chain must have an occupancy of exactly 1.0.
- The ligand chain must contain exactly one residue.
- The formula weight, according to the Chemical Component Dictionary (CCD), must fall within the range of [100, 900] Da.
- The CCD elemental composition is restricted to a subset of {H, C, O, N, P, S, F, Cl}.
- The ligand must possess at least 3 heavy (non-H) atoms.
- No covalent bonds are formed between the ligand and other chains in the complex.

##### Chemical Redundancy Removal

To evaluate the model’s generalization to structurally novel ligands, we identify a low-homology subset by excluding test ligand entities for which there exists a single training complex that contains both a protein with sequence identity ≥0.4 and a ligand with Tanimoto similarity ≥0.6 relative to the test case. This similarity is calculated using Morgan Fingerprints (radius 2, 2048 bits), ensuring the chemical diversity and novelty of the evaluation benchmark.

##### High-Quality Validation and Artifact Filtering

For ligand-polymer interfaces, we implement additional structural quality controls to eliminate potential artifacts:

- **RCSB Validation Metrics:** We retrieve instance-level validation data via GraphQL. A ligand is retained only if it exhibits an absence of intermolecular clashes and stereochemical outliers, maintains full atomic completeness (1.0), and satisfies thresholds for Real-Space *R*-factor (*RSR* ≤ 0.2) and Correlation Coefficient (*RSCC* ≥ 0.95). Only the instance designated as the “best instance” by the RCSB is used.
- **Crystallographic Artifact Removal:** To exclude binding poses stabilized by crystal packing rather than biological affinity, we perform a symmetry-mate contact check. We expand the unit cell into a 3 ×3 ×3 grid using the Biotite[11] library and identify neighbors within 5 Å via a KD-tree. Any ligand forming contacts with symmetry-related chains (i.e., chains outside the intersection of Assembly 1 and the asymmetric unit) is discarded.

### A.12 Antibody-Antigen Test Set Curation

To curate the antibody-antigen interface prediction benchmark, we utilize structural metadata from the Structural Antibody Database (SAbDab) [8]. We first establish a mapping from PDB identifiers and author-assigned chain IDs to specific antibody types, including single-chain variable fragments (*scFv*), heavy-light complexes (*HL*), and standalone heavy (*H*) or light (*L*) chains.

Based on this mapping, we implement an automated labeling protocol for the low-homology subset:

- **Individual Chains:** Any chain present in the sabdab database is annotated with its corresponding antibody type.
- **Complex Interfaces:** For structural interfaces, labels are assigned based on the composition of the interacting partners:
  - *Antibody-Antibody* : Interfaces where both participating chains are identified as antibodies.
  - *Antibody-Protein*: Interfaces involving one antibody chain and one non-antibody protein chain. These are further refined by the antibody component type, forming the core of our antigen-binding evaluation.

The Ab-Ag metrics we report are on the *Antibody-Protein* subset.

## Footnotes

1 https://github.com/bytedance/PXMeter

